# Extending Subcortical EEG Responses to Continuous Speech to the Sound-Field

**DOI:** 10.1101/2023.11.08.566173

**Authors:** Florine L. Bachmann, Joshua P. Kulasingham, Kasper Eskleund, Martin Enqvist, Emina Alickovic, Hamish Innes-Brown

**Affiliations:** Eriksholm Research Centre, Snekkersten, Denmark; Automatic Control, Department of Electrical Engineering, Linköping University, Sweden; Oticon A/S, Smørum, Denmark; Department of Health Technology, Technical University of Denmark, Lyngby, Denmark

**Keywords:** Electroencephalography, Temporal Response Function, Speech-ABR, Neural Speech Processing, Auditory Brainstem Response

## Abstract

The auditory brainstem response (ABR) is a valuable clinical tool for objective hearing assessment, which is conventionally detected by averaging neural responses to thousands of short stimuli. Progressing beyond these unnatural stimuli, brainstem responses to continuous speech presented via earphones have been recently detected using linear temporal response functions (TRFs). Here, we extend earlier studies by measuring subcortical responses to continuous speech presented in the sound-field, and assess the amount of data needed to estimate brainstem TRFs. Electroencephalography (EEG) was recorded from 24 normal hearing participants while they listened to clicks and stories presented via earphones and loudspeakers. Subcortical TRFs were computed after accounting for non-linear processing in the auditory periphery by either stimulus rectification or an auditory nerve model. Our results demonstrated that subcortical responses to continuous speech could be reliably measured in the sound-field. TRFs estimated using auditory nerve models outperformed simple rectification, and 16 minutes of data was sufficient for the TRFs of all participants to show clear wave V peaks for both earphones and sound-field stimuli. Subcortical TRFs to continuous speech were highly consistent in both earphone and sound-field conditions, and with click ABRs. However, sound-field TRFs required slightly more data (16 minutes) to achieve clear wave V peaks compared to earphone TRFs (12 minutes), possibly due to effects of room acoustics. By investigating subcortical responses to sound-field speech stimuli, this study lays the groundwork for bringing objective hearing assessment closer to real-life conditions, which may lead to improved hearing evaluations and smart hearing technologies.

## 1 Introduction

The Auditory Brainstem Response (ABR) is an electrophysiological measure of the subcortical neural activity of the auditory system in response to sound. It is widely used for hearing screening, especially in populations for which behavioral measures of hearing are not feasible, such as newborns (Galambos & Despland, 1980; Joint Committee on Infant Hearing, 2007) and coma patients (Guérit, 1999). The aggregate activity of several subcortical stages of the auditory pathway gives rise to the ABR event-related potential (ERP) which can be detected using electroencephalography (EEG) (Picton, 2010). Since the electric potentials generated by this neural activity are relatively weak when measured using scalp-EEG, conventional ERPs are computed using signal averaging methods, often with thousands of repetitive short stimuli, such as clicks (Picton, 2010), chirps (Rodrigues, Ramos, & Lewis, 2013), or short speech syllables (Chandrasekaran & Kraus, 2010). Although such methods provide robust ABRs, they are far removed from ecologically relevant stimuli, and recent years have seen a shift towards more complex stimuli relevant to daily life such as continuous speech in several fields of auditory and speech neuroscience (Brodbeck & Simon, 2020; Hamilton & Huth, 2020; Lunner et al., 2020).

Recent studies have found that subcortical responses to continuous natural speech can be reliably detected using deconvolution methods which produce a temporal response function (TRF) (Maddox & Lee, 2018). These subcortical TRFs show similar waveform morphologies to the conventional click ABR, with a prominent wave V peak, which is thought to arise from a mixture of subcortical structures, including the inferior colliculus (Møller & Jannetta, 1983; Moore, 1987; Starr, 1976). Although TRFs have been widely used in recent years to study *cortical* responses to continuous speech (Brodbeck & Simon, 2020; Di Liberto, O’Sullivan, & Lalor, 2015; Alickovic, Lunner, Gustafsson, & Ljung, 2019; Kulasingham et al., 2020), relatively few studies have investigated *subcortical* TRFs to continuous speech (Bachmann, MacDonald, & Hjortkjær, 2019, 2021; Maddox & Lee, 2018; Polonenko & Maddox, 2021; Shan, Cappelloni, & Maddox, 2023).

In accordance with conventional ABR measurements, these early studies computing subcortical responses to continuous speech involved stimuli presented via insert earphones. Although this allows good control over the exact stimulus presented at the ear, measurements of subcortical responses to speech in sound-field conditions could enable several other use-cases. Subcortical responses to speech presented in the sound-field could enable clinical tests of hearing impairment with natural conversational stimuli in a realistic setting, leading to increased levels of relevance and patient comfort. Critically, since hearing aids incorporate speech-specific signal processing and noise cancellation stages that may attenuate the short transient stimuli that are typically used for evoked ABR paradigms (Garnham, Cope, Durst, McCormick, & Mason, 2000), the use of sound-field speech stimuli could lead to more reliable measurements of subcortical responses from listeners wearing hearing aids. Indeed, sound-field speech stimuli have already been successfully used to measure *cortical* responses in hearing aid users, and the impact of various hearing aid settings on neural speech tracking (Alickovic et al., 2020, 2021; Carta, Alickovic, Zaar, Valdes, & Liberto, 2023).

Although conventional averaged ABRs using sound-field stimuli have been studied in animal models (Kim, Schrode, & Lauer, 2022; Land, Burghard, & Kral, 2016; Willott, 2006), there have been fewer studies in humans (Jarollahi et al., 2020; Schebsdat, Kohl, Corona-Strauss, Seidler, & Strauss, 2018). Sound-field Auditory Steady State Responses (ASSRs) have been more widely studied in humans (Hernández-Pérez & Torres-Fortuny, 2013; Stroebel, Swanepoel, & Groenewald, 2007; Zapata-Rodriguez, Laugesen, Jeong, Brunskog, & Harte, 2021), with a view towards similar applications in clinical diagnosis and hearing aid fitting (Damarla & Manjula, 2007; Shemesh, Attias, Magdoub, & Nageris, 2012). These studies highlight potential challenges when measuring sound-field responses, due to delays, reverberation, binaural interactions and other effects of room acoustics (Zapata-Rodríguez, 2020). Considering that the TRF is estimated using a predictor based on the stimulus, distortions of the stimuli that reach the ears due to propagation through the sound field might either result in a distorted TRF estimate or increase the amount of data needed to estimate the TRF. The latter may be partly counteracted by employing predictors generated using an auditory nerve model (Zilany, Bruce, Nelson, & Carney, 2009; Zilany, Bruce, & Carney, 2014) instead of simple rectification, which has shown to substantially improve estimated TRFs (Shan et al., 2023; Kulasingham et al., 2023). Still, it remains unclear whether subcortical TRFs, which require temporally precise measurements, can be detected using continuous speech in the sound-field.

In this work, we investigated subcortical EEG responses to continuous speech in both insert earphone and sound-field conditions. We analyzed EEG data collected from 24 participants with normal hearing and used TRF methods to estimate subcortical responses. First, we explored morphological differences in subcortical responses to sound presented in the sound-field versus via insert earphones in the same participants, for both subcortical TRFs to speech stimuli and conventional click-ABRs. Specifically, we examined whether after accounting for the speaker delay, the wave V peak in the sound-field condition occured at similar latencies as in the insert earphone condition (Research Question 1). Second, we compared subcortical TRFs estimated using predictors generated using simple rectification or an auditory nerve model (ANM) (Zilany et al., 2009, 2014) (RQ2) and validated prior work showing that the ANM substantially improves estimation of wave V peaks. Third, we studied whether distortions of the stimulus in the sound-field leads to a need for more data (longer recording time) to estimate TRFs with response peaks above the noise floor (RQ3). Finally, we investigated whether subcortical responses to speech in the sound-field could be detected in all participants (RQ4).

Our work may lead to reliable objective measures of subcortical auditory activity using ecologically relevant stimuli presented in the sound-field, and lays the groundwork for future investigations into utilizing subcortical responses for clinical evaluations of hearing impairment and hearing aid fitting in more realistic environments.

## 2 Materials and methods

### 2.1 Experimental setup

#### 2.1.1 Procedure

EEG data were collected from 24 participants (14 males, *M*_*age*_=37.07, *SD*_*age*_=10.02 years) with clinically normal and symmetric hearing (all pure-tone thresholds in octave steps from 250 to 4000 Hz *≤* 20 dB HL, no more than 15 dB HL difference between ears at each frequency). All participants were native Danish speakers, right-handed, and provided written informed consent. The study was approved by the ethics committee for the capital region of Denmark (journal number 22010204). The experiment consisted of several parts for the insert earphone and sound-field conditions, and it was balanced whether participants started with the inserts or the sound-field condition. Story order was pseudo-randomized, presenting stories in the same order to two age-matched participants with different starting conditions. The insert earphone condition consisted of the following stimuli: speech (47 minutes 56 seconds, divided in 8 audiobook segments), clicks (5 minutes), control condition with clicks and insert eartips outside the ears (5 minutes). The sound-field condition consisted of the same speech (47 minutes 56 seconds) and click material (5 minutes).

#### 2.1.2 Stimuli

EEG data were collected while participants listened to clicks and four stories from adventures by H.C. Anderson read by a male speaker. The five minutes of click trains were presented both via insert earphones and in the sound-field. Click trains consisted of 44.1 clicks of alternating polarity per second. The presentation time of each click was randomly distributed according to a pseudo-Poisson distribution, limiting the minimum inter-click-interval to 15ms. Clicks were rectangular and of 91 µs duration (4 samples, the closest approximation to 100 µs at a 44.1 kHz sampling rate). Click stimuli were presented at 72 dB peak-to-peak equivalent SPL, matching their peak-to-peak voltage to that of a 1 kHz pure-tone presented at 72 dB SPL.

Stories were divided into two segments, resulting in 8 trials (*M*_*duration*_ = 6 minutes 0 seconds, *SD*_*duration*_ = 55 seconds). All trials were presented in two conditions, through insert earphones or through a loudspeaker in the sound-field. Before presentation, the original 2-channel audiobook with a sampling frequency of 44.1 kHz was averaged to construct a mono signal, and high-pass filtered at 1 kHz using a 1^*st*^ order Butterworth filter because the neural response from the brainstem is mainly driven by high frequencies (Abdala & Folsom, 1995). Each of the 8 trials were scaled to have the same root-mean-square (RMS) value as a 1 kHz pure tone at 72 dB sound pressure level (SPL).

Participants were seated facing the loudspeaker at ear height and a distance of 1.5 m (approximately 4.4 ms sound travelling delay) and instructed to relax and listen to the audio, while looking at a fixation cross in front of them. The room was quiet and insulated from outside noise, with dimensions of 4.5*×*3.2 *×*2.5 meters and a reverberation time (*RT*_60_ estimated with *T*_30_) of 0.32 seconds. All audio was replayed with a sampling rate of 44.1 kHz using an RME Fireface UCX audio interface (RME Audio, Haimhausen, Germany). In the insert condition, sound was delivered via Etymotic ER-2 (Etymotic Research, Illinois, USA) insert earphones (approximately 0.9 ms sound travelling delay), which were shielded using a grounded metal box to avoid direct stimulus artifacts on the EEG. Additionally, a control condition where the clicks were played while the insert earphones were not placed in the ear (but with the earphone wires in the same locations near the participant) was recorded to check whether there were electrical stimulus artifacts in the EEG. Visual inspection confirmed that stimulus artifacts were not present in the ERPs estimated from the control condition, or in any other conditions. For the sound-field condition, the stimuli were presented using a Genelec 8040A loudspeaker (Genelec Oy, Iisalmi, Finland).

### 2.2 EEG preprocessing

EEG data were collected using a Biosemi Active 2 system (BioSemi, Amsterdam, The Netherlands) at a sampling frequency of 16,384 Hz. A 32-channel cap with electrodes in standard 10–20 positions was used. In addition, electrodes were placed on the left and right mastoids, left and right earlobes and above and below the right eye. Data analysis was conducted in MATLAB (version R2021a) and the Eelbrain Python toolbox (version 0.38.1) (Brodbeck et al., 2021). Only the Cz channel was used for further analysis, referenced to the average of the two mastoid channels. A 1^st^ order Butterworth highpass filter with a cutoff frequency of 1 Hz was applied. Next, to remove power line noise, FIR notch filters with widths of 5 Hz and center frequencies at all multiples of 50 Hz up to 1000 Hz were applied. High amplitude sections of the EEG data that were 5 standard deviations (SD) above the mean were assumed to be contaminated with artifacts, and 1 second segments around these points were set to zero in both the EEG data and the predictors used for later TRF analysis (mean percentage of data set to zero = 2.9%, SD = 2.7%). Only the data from 2 to 242 seconds in each of the 8 trials was used for further analysis to avoid possible onset and offset artifacts or responses.

Since subcortical responses consist of peaks in the waveform that are only a few milliseconds in duration, it is crucial that the EEG data and the presented stimuli are synchronized with millisecond precision. This synchronization could be impacted by timing jitters in the trigger or clock drifts between the stimulus presentation system and the EEG recording system. To avoid these issues and ensure synchronization, the audio output of the RME soundcard was fed into the Biosemi system as an external sensor on the Erg1 channel via an optical isolator (StimTrak, BrainProducts, GmbH, Gilching, Germany) to maintain electrical separation. The recorded signal on the Erg1 channel of the EEG system was subsequently used to detect click onsets for ERPs and to generate speech predictors for TRF analysis.

### 2.3 Click ERP Estimation

The click evoked ABR was estimated on EEG data that was further filtered between 30 and 1000 Hz using a 1^*st*^ order Butterworth IIR bandpass filter. Click onsets were detected using the Erg1 channel and EEG epochs -10 to 30 ms around the click onset were extracted. This resulted in 13185 epochs that were averaged to estimate the ERP for each condition (inserts, sound-field and control conditions). The standard procedure of averaging both condensation and rarefaction click responses together was followed. The ERPs were then baseline corrected by subtracting the average of the pre-stimulus activity from -10 to 0 ms.

### 2.4 Speech Predictors

The speech stimuli were used to construct two types of predictors for the TRF model: Rectified Speech (RS) predictor and Auditory Nerve Model (ANM) predictor. These predictors approximate the processing stages of the peripheral auditory system, and thereby account for non-linearities that are hard to fit using linear TRF models. Since the brainstem response is largely unaffected by the polarity of the auditory input, two predictors were estimated for each case using the speech signal recorded on the Erg1 channel and its sign-flipped (inverted) version. The RS predictors were constructed by rectifying the speech signals, leading to predictors that only kept either the positive or the negative peaks, in accordance with prior work (Maddox & Lee, 2018). This procedure serves as a very coarse approximation of the rectification properties of the peripheral auditory system.

The ANM predictor was formed using a model of the auditory periphery (Zilany et al., 2014), in line with recent work that showed this method improved TRF estimates (Shan et al., 2023; Kulasingham et al., 2023). In brief, the speech stimuli (and its polarity inverted version) were used as inputs to this model, which modelled 43 high spontaneous rate auditory nerve fibers with center frequencies logarithmically spaced between 125 Hz and 16 kHz. The outputs of this model were 43 mean firing rate signals, which were then averaged to form the final predictor pair (i.e., for the original speech signal and its polarity inverted version). The implementation in the cochlea python toolbox (Rudnicki, Schoppe, Isik, Völk, & Hemmert, 2015) (https://github.com/mrkrd/cochlea) was used to generate these predictors. To ensure that the estimated TRF peak latencies were not affected by inherent lags in the auditory model, the ANM predictors were delayed by 1.1 ms in order to have the maximum correlation with the RS predictor. For further comments on the suitability of this method of accounting for inherent lags in the auditory model, please see the Discussion.

### 2.5 Temporal Response Function Estimation

The RS and ANM predictors were used to estimate TRFs for each experimental condition. The TRF is a linear model that represents the time-locked activity of the neural system to the given predictor. The frequency domain method given in previous studies (Maddox & Lee, 2018; Polonenko & Maddox, 2021) was used to estimate the TRF:

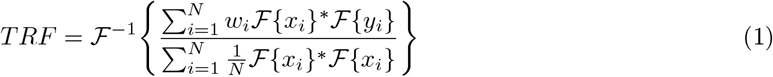

Here, ℱ denotes the Fourier transform, *N* is the number of trials, *x*_*i*_, *y*_*i*_ and *w*_*i*_ are the predictor, EEG signal and weight for trial *i*, and ^*∗*^ denotes the complex conjugate. The trial weights *w*_*i*_ were set to be the reciprocal of the variance of the EEG data of trial *i* normalized to sum to 1 across trials, in line with prior work (Polonenko & Maddox, 2021). This was done to down-weight noisy (high variance) EEG trials. The resulting TRF has lags ranging from *− T/*2 to *T/*2 where *T* is the data length.

Two TRFs were estimated separately for each predictor pair (i.e, generated from the speech signal and its polarity-inverted version), and then averaged together. These TRFs were then bandpass filtered between 30-1000 Hz using a delay compensated FIR filter and smoothed using a Hamming window of width 2 ms. This smoothing step helped to eliminate high frequency noise in the TRF and led to cleaner wave V estimates. After this, the TRF segment from -10 to 30 ms was extracted for further analysis and the mean baseline activity from -10 to 0 ms was subtracted from each TRF. Finally, the TRFs were scaled to have the same RMS as the click ERPs averaged across all participants, for enabling morphology comparison with the click ERPs.

To investigate the effect of data length on TRF estimation, TRFs were fit on a consecutively increasing number of trials (i.e, 2, 3, …, 8 trials, corresponding to 8, 12, …, 32 minutes of data) in the order that they were presented in the experiment. This simulates TRF estimation as if the experiment had been terminated after a variable number of trials. For each data length, a leave-one-out cross-validation approach was followed, with one trial being used as test data to estimate model fits and the other trials being used to fit the TRF. The TRFs for each cross-validation fold were averaged together to form the final TRF for that data length. A null model was formed by averaging the TRFs that were fit on circularly shifted predictors (shifts of 30, 60 and 90 seconds), similar to typical null models used in prior work with cortical TRFs (Kulasingham et al., 2020). This method preserved the temporal characteristics of the predictor, while destroying the alignment between the predictor and the EEG signals. The same leave-one-out cross-validation approach at each data length was followed for the null models.

### 2.6 Performance metrics

The Pearson correlation between the predicted EEG from the TRF model and the actual EEG was used as an estimate of the model fit, in line with previous TRF studies (Crosse, Di Liberto, Bednar, & Lalor, 2016; Kulasingham & Simon, 2023). The TRF estimated on the training trials was used to predict the EEG on the test trial using the leave-one-out procedure, and the average of the prediction correlations across folds was used as the model fit. The null model fits were also estimated in the same manner using the null model TRFs.

The wave V peak of the subcortical TRFs was extracted by detecting the largest peak between 4-9 ms. The SNR of the peak amplitude was calculated as SNR = 10 log_10_(*S/N*) where the power in a 5 ms window around the peak was used for the signal power *S* and the noise power *N* was estimated as the average TRF power in 5 ms windows in the range -500 to -20 ms. The SNR was set to have a minimum value of 0 dB, since negative SNRs (signal below the noise floor) lack meaningful interpretation. The threshold for a meaningful wave V peak was set at 3 dB (signal power is twice the noise power), since this SNR corresponded with visually distinct peaks.

### 2.7 Statistical Analysis

The model fits for the full data length ANM and RS TRFs in the inserts and sound-field conditions were compared using related measures two-tailed t-tests with Holm-Bonferroni multiple comparisons correction. The null model fits for each participant were subtracted before the comparison. Cohen’s d effect sizes, *t*-values and *p*-values are reported. For testing differences in wave V SNR, non-parametric statistical tests were employed, since the wave V SNRs had a skewed distribution with participants without clear wave V peaks having 0 dB SNR. Wave V SNRs for the ANM and RS TRFs at the full data length were compared between selected conditions using paired two-tailed small sample Wilcoxon signed rank tests with Holm-Bonferroni multiple comparisons corrections. Next, the same tests were used to examine differences in ANM TRF wave V SNR between insert earphone vs. soundfield conditions at different data lengths. The group medians, test statistics (rank sums above zero) and *p*-values are reported. Two participants were excluded from the statistical tests since they did not have data for the full 32 minutes.

## 3 Results

### 3.1 Subcortical responses for speech presented in the sound-field versus through insert earphones

The EEG data from the midline central (Cz) channel referenced to the average of the two mastoid channels was used to estimate click evoked ERPs and subcortical TRFs to continuous speech. The TRFs were estimated using either the rectified speech (RS) or the auditory nerve model (ANM) predictors. The average click ERPs and speech TRFs for both speech presented via insert earphone and in the sound-field are shown in Fig. 2 (top). Clear wave V peaks were seen on group-level for both the click ERPs and the speech TRFs. The pre-stimulus period (−10 to 0 ms) for the click ERP shows activity from the previous click response, since the click train had inter-click spacing mostly ranging from 15-25 ms. Even though the inter-click latency was randomly jittered (see Methods) there is still a smeared response to the previous click in Fig. 2. However, this does not affect any further conclusions since the click ERP was investigated merely as a reference stimulus for inserts and speaker conditions, and the wave V peaks are still clearly visible in both cases at the appropriate latencies within 5-10 ms. The fact that the earlier waves I-IV are not visible may be due to the fast click rate, which could lead to weaker early waves (Picton, 2010).

**Figure 1:**
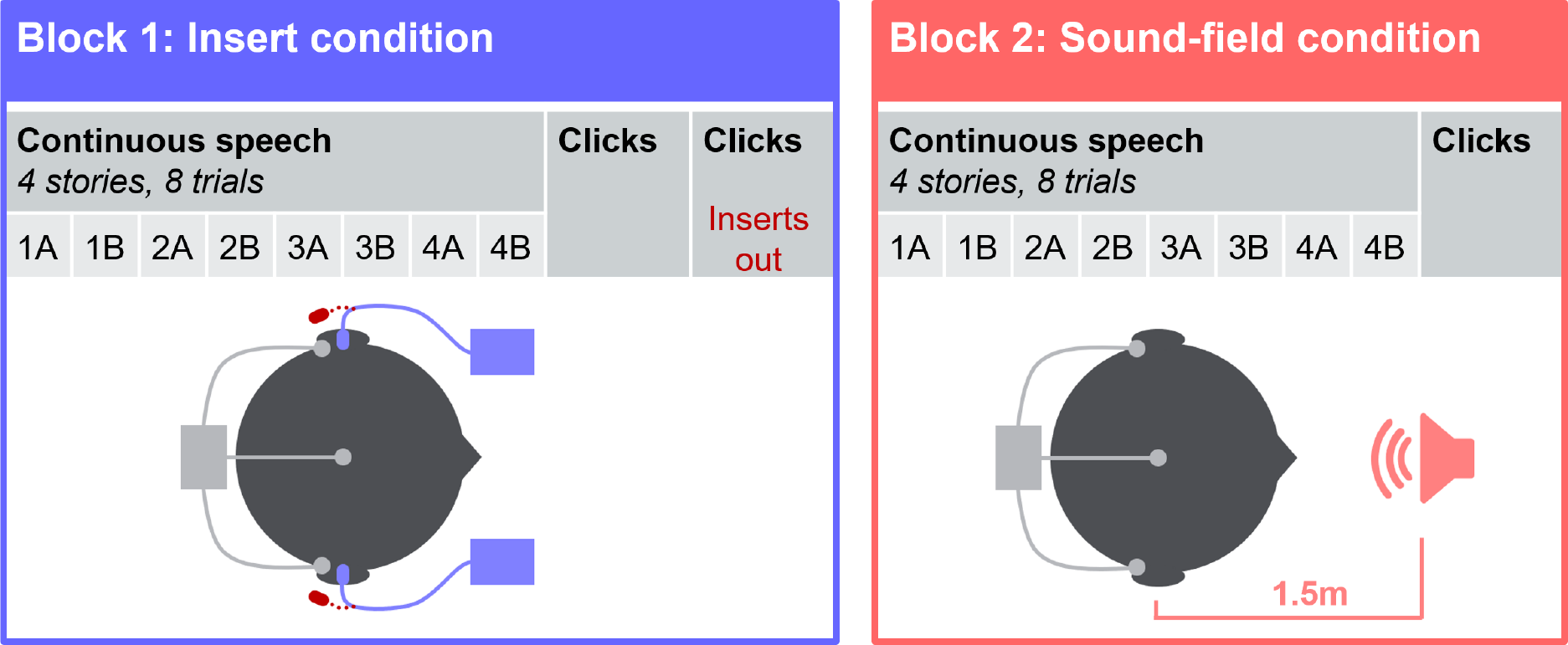
Experiment overview. Continuous speech and click stimuli were presented in an insert earphone condition block, and a sound-field condition block, balancing the starting condition across participants. The speech consisted of four stories, each split into two parts (A and B). Story presentation order was pseudo-randomized across participants and identical across conditions for a given participant. In the insert condition, clicks were also presented with earphone tips outside the ear, serving as a control condition.

**Figure 2:**
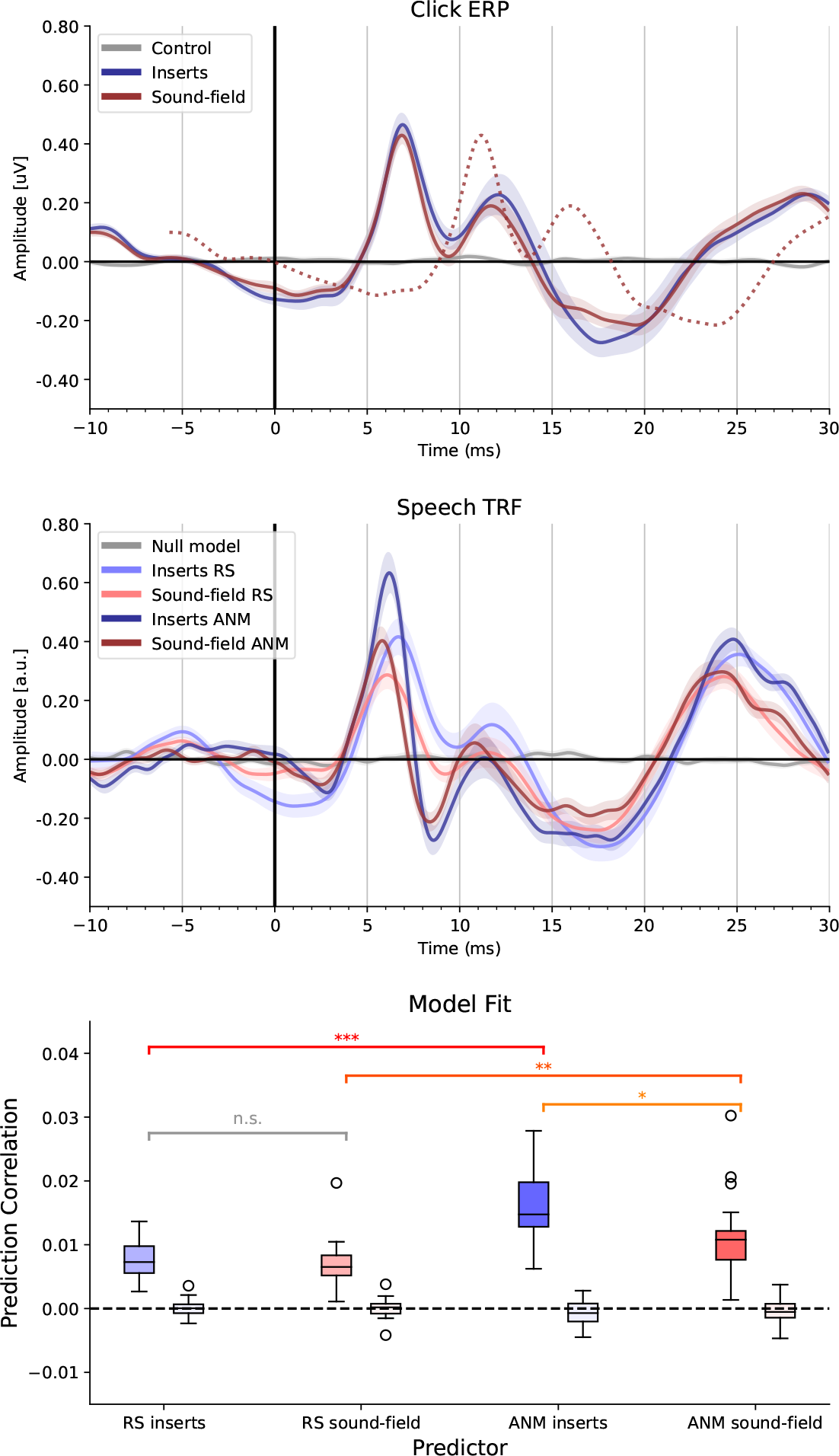
Group-level click ERPs and speech TRFs in inserts and speaker conditions. **Top**: The group-level click ERP across 24 participants is shown for both inserts (blue) and sound-field conditions corrected (red solid) and uncorrected (red dotted) for speaker delay of 4.4 ms. A clear wave V peak is seen that is consistent for both conditions. The pre-stimulus activity and lack of earlier waves I-IV may be due to the fast repetition rate of the click stimuli. The control condition with insert eartips outside the ears is depicted in grey and has very low amplitude. **Middle**: The group-level speech TRF across 24 participants is shown for both inserts and speaker conditions for the RS and ANM predictors. Clear wave Vs are seen in all cases, but the ANM predictor results in narrower and larger wave Vs. The amplitude of the wave V is reduced for all speaker conditions, possibly due to effects of room acoustics. The null model is also shown in grey for visual comparison with the noise floor. **Bottom**: The model fit prediction correlations for the estimated TRFs. Boxplots are shown across participants for each condition and predictor, and the null model fits are indicated by the lighter colored boxes next to them. All TRF models significantly outperform the null models. The ANM TRF model fits are significantly higher than the RS TRF model fits in both conditions (* *p <* 0.05, ** *p <* 0.01, *** *p <* 0.001).

The general response morphologies across sound-field and insert earphone condition showed close resemblance. For illustrative purposes, the sound-field response without correcting for the travelling time of sound from the speaker to the eardrum (4.4 ms) is also indicated for the click stimulus (dashed line in Fig 2 top). Henceforth, all results will be reported after accounting for this delay, as well as an earphone presentation delay of 0.9 ms. After accounting for both these sound travelling delays, the average latencies of wave V peaks were similar across inserts and sound-field conditions for both click ERPs (mean [SD] of click ERP wave V latency for inserts = 7.0 [0.4] ms, sound-field = 6.9 [0.5] ms) and speech TRFs (RS inserts = 6.6 [0.6] ms, RS sound-field = 6.2 [0.6], ANM inserts = 6.4 [0.4], ANM sound-field = 6.0 [0.4]). As in previous work, response peaks for the speech occurred slightly earlier than for clicks (Maddox & Lee, 2018). This difference was more pronounced in the sound-field than the insert earphone condition, which will be addressed in the Discussion. The sound-field conditions resulted in lower amplitudes compared to the insert condition for all cases (percentage reduction in average wave Vpeak amplitude for sound-field compared to inserts: click ERP = 9.2%, speech TRF RS = 31.4%, speech TRF ANM = 34.7%). This could be due to differences between the presented stimuli and the actual signal at the ear for the sound-field condition, due to room acoustics and reverberation. This mismatch could lead to a smearing of the wave V peak and hence a lower amplitude. A more thorough exploration of possible issues with subcortical responses to sound-field stimuli is provided in the Discussion.

### 3.2 Responses computed to predictors obtained using simple rectification versus an auditory nerve model

The average wave V latency for the ANM predictor was slightly earlier than that for the RS predictor, even though the model delays in the ANM predictor were compensated for by shifting the ANM predictor to have the maximum correlation with the RS predictor (see Discussion). Qualitatively, responses obtained with the RS predictor show a broader peak than responses computed with the ANM predictor. The amplitudes of the TRF wave V were scaled to be comparable to the amplitudes of the click ERP wave V using a common scaling factor for all participants. The individual speech TRFs for the ANM predictor also show clear wave V peaks and consistent TRF waveforms for all participants (see Fig. 4).

The model fits of the estimated speech TRF models are shown in the bottom row of Fig 2. All the estimated model fits are well above the null model fits. The model fits for each case were compared after subtracting the individual null model fits using paired *t*-tests with Holm-Bonferroni correction. The ANM model fits were significantly higher than the RS model fits for both inserts (*d* = 1.79, *t*_23_ = 8.19, *p <* 0.001) and sound-field conditions (*d* = 0.91, *t*_23_ = 4.19, *p* = 0.0013). The RS TRF model fits were not significantly different across inserts and sound-field conditions (*d* = 0.097, *t*_23_ = 0.442, *p* = 0.66). However, the ANM TRF model fits were significantly higher in the inserts compared to the sound-field condition (*d* = 0.67, *t*_23_ = 3.08, *p* = 0.011). These results indicate that the ANM leads to a better TRF estimate than the RS, and that the insert condition leads to better estimates than in the sound-field, possibly due to effects of room acoustics.

### 3.3 Amount of data needed for subcortical responses to continuous speech

Next, we investigated the amount of data required to estimate reliable TRFs using two metrics; the model fit prediction correlations and the wave V SNR. The latter is a measure of the wave V amplitude relative to the noise floor (see Methods), and has been previously used to evaluate subcortical TRFs (Polonenko & Maddox, 2021; Shan et al., 2023). TRFs were fit on a sequentially increasing number of 4 minute trials, and both model fit and wave V SNR were calculated (see Fig. 3). The ANM predictor outperformed the RS predictor in all cases. The sound-field condition had lower wave V SNR than the inserts conditions, in line with the reduced peak amplitudes seen in the TRFs. ANM TRF model fits and wave V SNRs for the insert condition were above the noise floor for all participants with only 12 minutes of data. For ANM TRFs when speech was presented in the sound-field, model fits were above zero in all participants with 12 minutes of data, too, whereas wave V SNR was 3 dB above the noise floor for all participants after an EEG recording time of 16 minutes and more.

**Figure 3:**
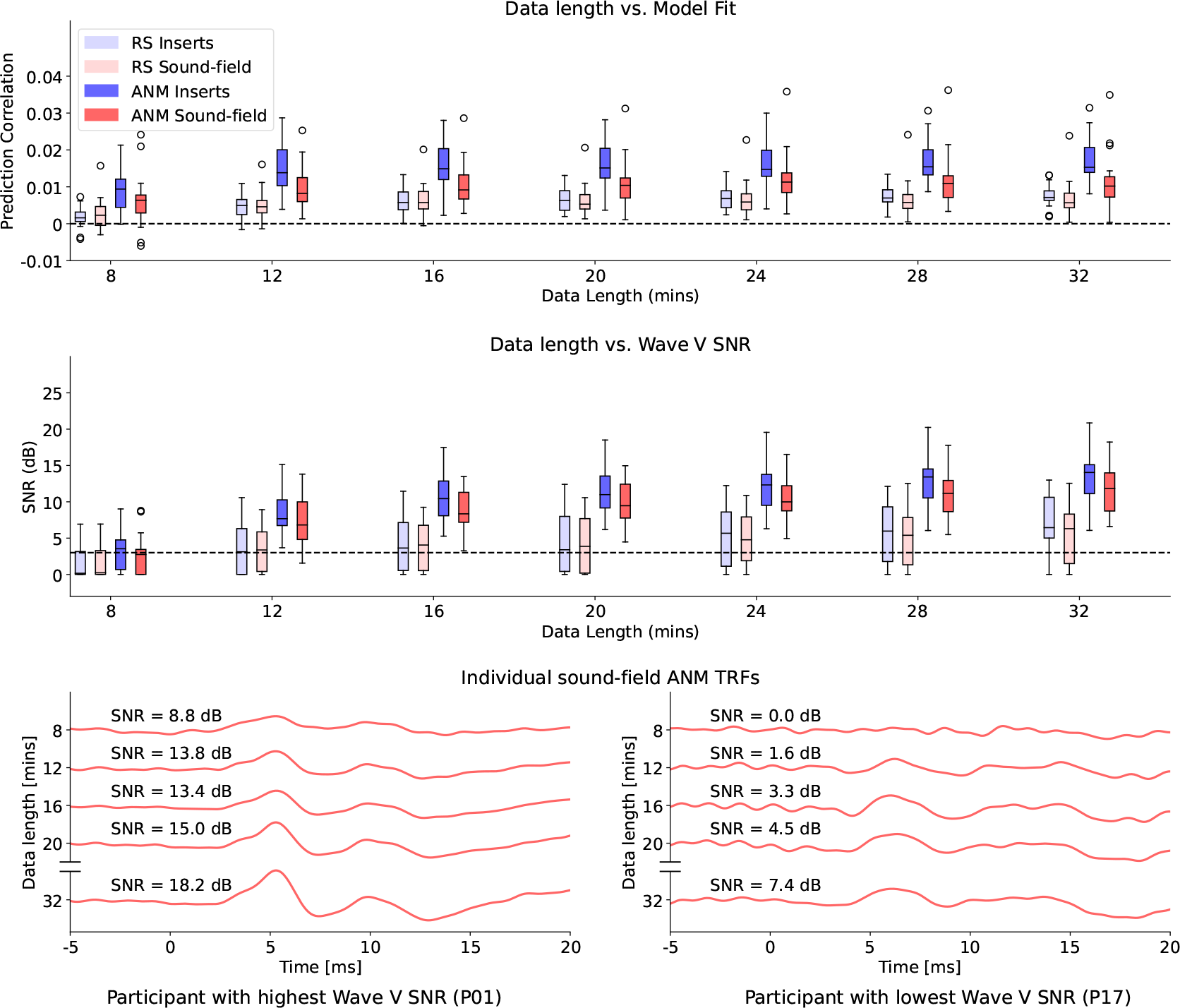
Impact of data length and predictor type. All boxplots are shown across participants. **Top**: Impact of data length on model fit prediction correlations. **Middle**: Impact of data length on wave V SNR. There is a clear increase in both model fits and wave V SNRs with increasing data length. The ANM model outperforms the RS model in all conditions, and the inserts condition seems to have both larger model fits and wave V SNRs than the sound-field condition. Interestingly, ANM TRFs with model fits above zero and wave V SNR above 3 dB for all participants were obtained with only 12 and 16 minutes of data for inserts and sound-field conditions, respectively. **Bottom**: Representative sound-field ANM TRFs of different data lengths for two participants.

**Figure 4:**
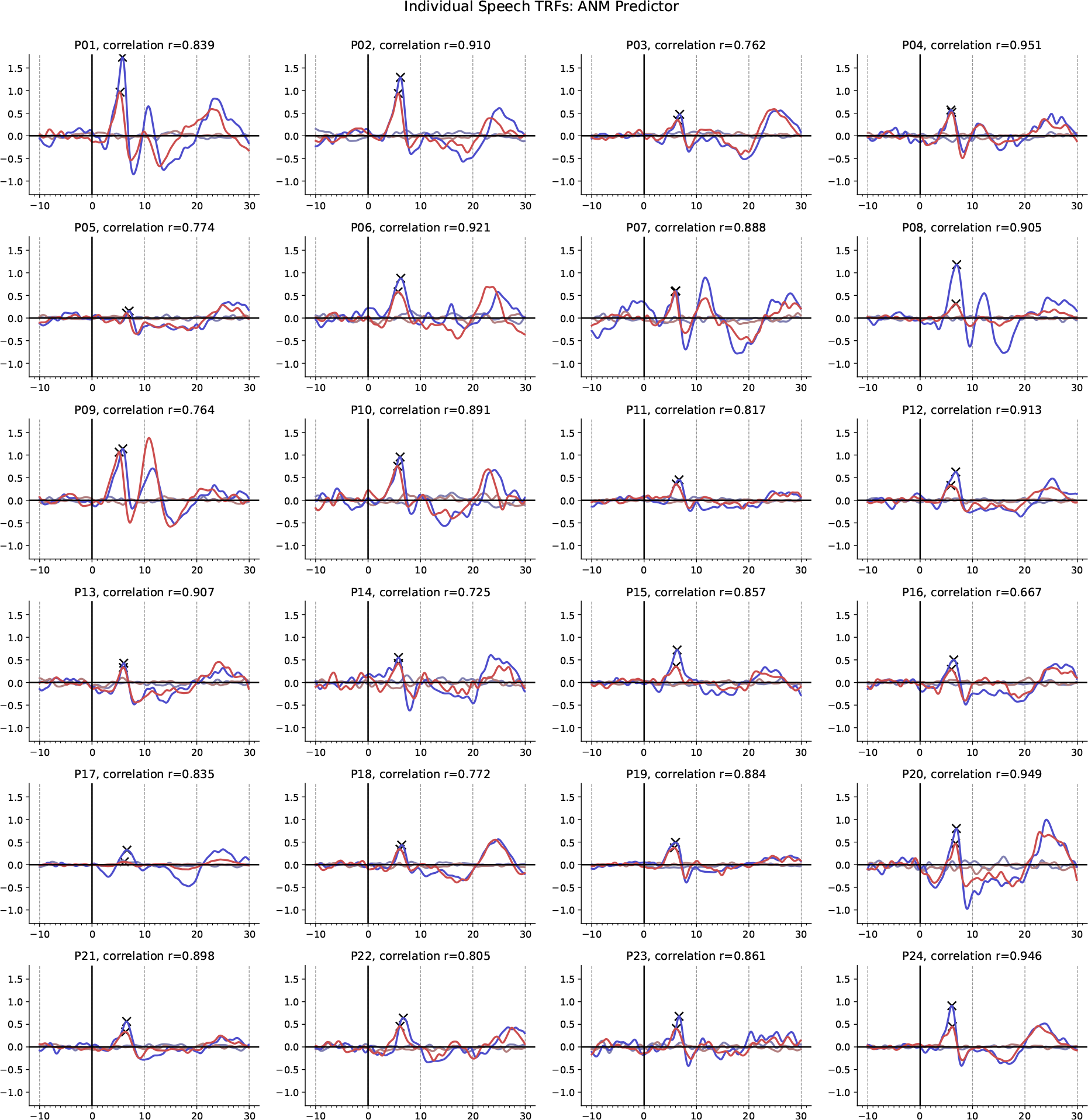
Individual speech TRFs for the ANM predictor. Individual participant TRFs are shown for both insert earphone (blue) and sound-field (red) conditions, along with corresponding null models (lighter colors). Clear wave V peaks are obtained in all participants and the TRFs for both conditions are similar on a single participant level (Pearson correlation between the TRF waveforms of both conditions are shown on top of each subplot).

To further investigate these trends, we used non-parametric pairwise Wilcoxon signed rank tests with Holm-Bonferroni multiple comparisons corrections on the wave V SNRs (see Methods). The ANM TRF SNRs were significantly higher than the RS TRF SNRs for the full 32 minutes of data (inserts RS median = 6.55 dB, ANM median = 14 dB, *T* = 0, *p <* 0.001; sound-field RS median = 6.32 dB, ANM median = 11.97 dB, *T* = 1, *p <* 0.001). We next restricted our statistical tests to only the ANM TRF SNRs and investigated the difference between the insert earphone SNRs and the sound-field SNRs. The tests were not conducted on 8 minutes of data, since several participants did not have clear wave V peaks (0 dB SNR). For 12 minutes of data, all participants were above 3 dB for the insert earphone condition, and the SNRs for the insert earphone condition (median = 8.29 dB) were significantly higher (*T* = 53, *p* = 0.031) than the SNRs for the sound-field condition (median = 7.23 dB). This indicated that the sound-field TRFs did not have wave V peaks that were as clear as those for the insert TRFs. We then investigated the amount of data required to reach similar SNRs for the sound-field TRFs. There was no significant difference between the sound-field TRF SNRs at 16 minutes (median = 8.47 dB) and the insert TRF SNRs at 12 minutes (*T* = 106, *p* = 0.524). Furthermore, the sound-field TRF SNRs at 20 minutes (median = 9.88 dB) were significantly higher than insert TRF SNRs at 12 minutes (*T* = 44, *p* = 0.018). These tests indicate that the sound-field TRF wave Vs are comparable to the insert earphone TRF wave Vs after an additional 4 minutes of data.

### 3.4 Consistency of individual responses to clicks and speech in insert and sound-field conditions

Finally, the individual responses as well as the consistency of wave V peaks across individuals was investigated. Clear wave V peaks could be seen for all participants for the click ERPs and the speech ANM TRFs (see Fig. 4 and Supplementary Figures S1, S2 for individual RS TRFs and click ERPs). As discussed above, average wave V latencies and amplitudes show some differences across insert and soundfield conditions (see Fig. 2), possibly due to room acoustics. However, these effects should primarily result in a systematic shift and the distribution of individual responses should be largely consistent, as evidenced by the individual TRFs in Fig. 4. To investigate this, we correlated individual peak V amplitudes and latencies across insert and sound-field conditions for both click ERPs and speech TRFs (see Fig. 5). The resulting Pearson correlation values were significant in all cases after correcting for multiple comparisons using the Holm-Bonferroni method (correlations and *p*-values are provided on Fig. 5 titles). This consistency could be due to a variety of individual factors such as signal quality, anatomical details (e.g., thickness of the scalp leading to larger responses for some participants), and properties of the auditory system. Although further investigation is required to characterize systematic differences such as the reduced amplitude in sound-field conditions, it is clear that subcortical responses can be reliably detected in sound-field conditions that are mostly consistent with responses detected using insert earphones on an individual basis.

**Figure 5:**
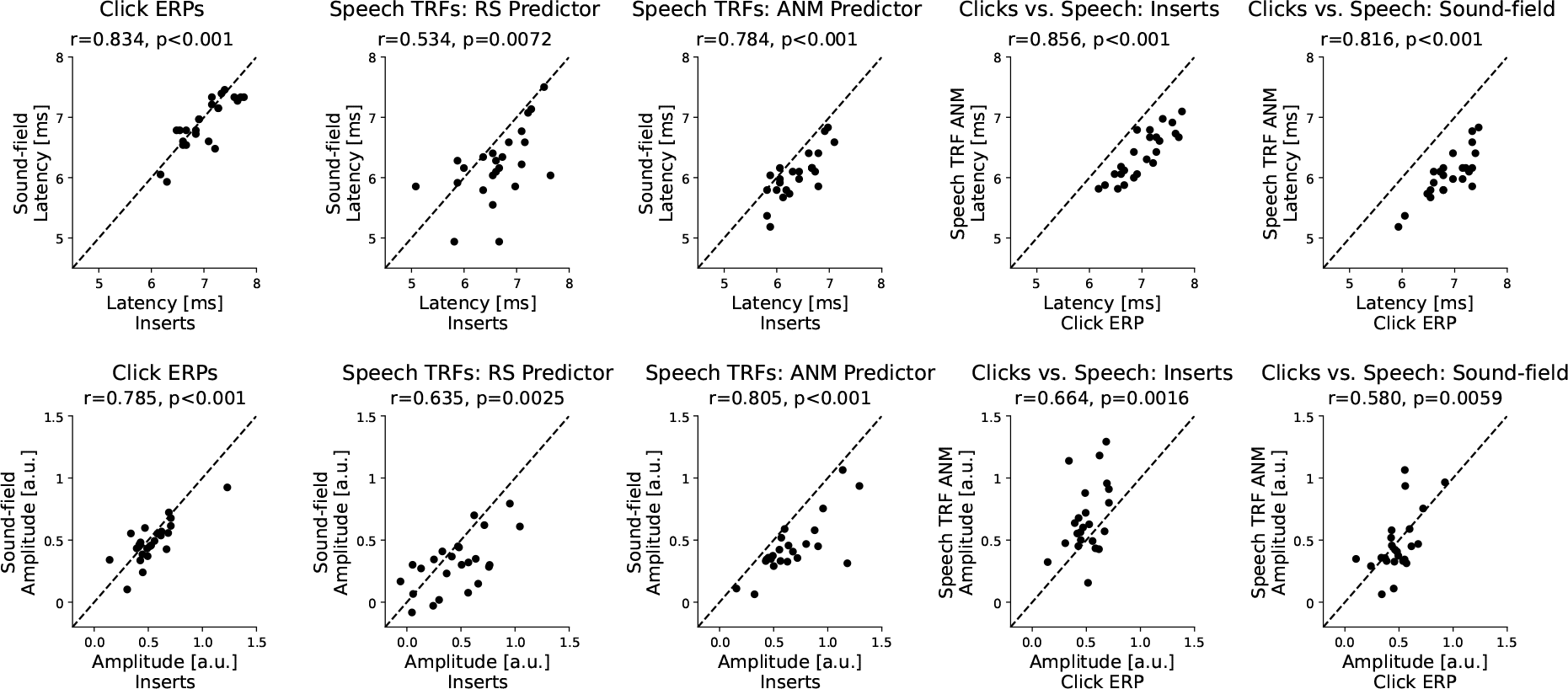
Individual wave V peak latencies and amplitudes. Scatterplots across participants are shown. **Top row**: wave V peak latencies, **Bottom row**: wave V peak amplitudes. The 3 leftmost columns show the comparison between inserts and sound-field for click ERPs, Speech TRF with RS predictor and Speech TRFs with the ANM predictor. The Pearson correlations are shown in each title, along with the corresponding p-value (Holm-Bonferroni corrected). The wave Vs in the insert and sound-field show a high degree of correlation, especially for click ERPs and ANM TRFs. The 2 rightmost columns show the comparison between click ERPs and speech ANM TRFs for inserts and sound-field. There is a high degree of correlation for both conditions, indicating that click ERPs and speech ANM TRFs have similar wave Vs. The speech ANM TRF shows an earlier wave V latency compared to the click ERP for all participants (see Discussion).

## 4 Discussion

### 4.1 Subcortical TRFs show clear wave V peaks in both insert earphone and sound-field conditions

Brainstem responses to clicks and speech were detected using ERPs and TRFs respectively, with the most prominent feature being the wave V peak. Speech TRFs were estimated using a simple rectified speech predictor (RS) and a complex auditory nerve model (ANM) predictor. The wave V was detected both when sound was presented through insert earphones and in the sound-field for click ERPs, RS TRFs and ANM TRFs. After accounting for sound propagation delays in the sound-field, response latencies for TRFs generated with the different predictors showed high similarity. TRF amplitudes were reduced when speech was presented in the sound-field versus via insert earphones. TRF estimations using an auditory nerve model consistently outperformed those obtained with simple rectification. Employing the auditory nerve model, slightly more data was required to obtain significant wave V responses from all participants when speech was presented in the sound-field (16 minutes) versus via insert earphones (12 minutes). Clear individual subcortical responses to continuous speech presented in the sound-field were observed for each individual participant.

### 4.2 Challenges when presenting stimuli in the sound-field

This experiment was conducted in a quiet room, though it was not anechoic or completely sound-proof. Distortions in the signal at the eardrum compared to the stimulus waveform may have occurred due to distortions in the transducers used, reverberation, binaural interactions, and other factors of room acoustics in the sound-field condition. Prior work has shown that room acoustics such as reverberations detrimentally impact both subcortical responses such as ASSRs (Zapata-Rodríguez, 2020) and behavioral measures such as speech intelligibility (Hodgson & Nosal, 2002). The effects of room acoustics lead to a mismatch between the heard signal and the stimulus used to generate the predictor. This may result in a ‘noisy’ linear model which may explain the reduced peak amplitudes seen in our sound-field TRFs. Differences in peak latency should mostly be captured by accounting for the sound travelling delay from the loudspeaker. Indeed, our results confirm that the wave V latencies were quite consistent across both insert and sound-field conditions once the propagation delay was accounted for. However, the observation that responses to speech occurred earlier than to clicks, as previously described by (Maddox & Lee, 2018), was more pronounced in the sound-field (RS: 0.7 ms, ANM: 0.9 ms) compared to the insert earphone (RS: 0.4 ms, ANM: 0.6 ms) condition. Effects of room acoustics might also play a role here. Reverberation time in a given room varies for different frequency bands (Zapata-Rodriguez et al., 2021) and might thus differently affect the click and speech stimuli with their respective frequency content. This may be one underlying reason how effects of room acoustics also have a distinct influence on perceived loudness. Stimuli calibrated to the same intensity at the eardrum are perceived louder when presented through loudspeakers than with headphones, an effect which is more pronounced for lower frequencies (Denk, Kohnen, Llorca-Bofí, Vorländer, & Kollmeier, 2021). As louder perceived stimuli show shorter ABR wave V latencies (Serpanos, O’Malley, & Gravel, 1997) and speech contains more energy at lower frequencies than the click sounds, this may lead to earlier ABR wave V peaks in the sound-field, especially for speech stimuli. The impact of room acoustics should thus be considered when using sound-field stimuli for clinical applications, perhaps by modelling the room acoustics and convolving the stimuli with the room impulse response (Zapata-Rodriguez et al., 2021) before generating the predictor. Another option would be to generate predictors from a recording of the audio signal heard at the ear, which may be suitable for applications using hearing aids with built-in microphones. However, our work shows that even without applying these corrections, it is possible to detect meaningful subcortical responses to sound-field speech stimuli, thereby laying the groundwork for applications of such techniques in settings that may not be as tightly controlled.

### 4.3 Predictors derived from auditory models improve TRF estimates

Consistent with prior work (Shan et al., 2023; Kulasingham et al., 2023), we show that the ANM predictor greatly outperforms the RS predictor in terms of model fits, wave V SNRs, and amplitude of wave V peaks at an individual participant level. However, it should be noted that the ANM predictor also resulted in slightly earlier wave V peaks (around 0.2 ms) compared to both the RS predictor and the click ERPs. This difference in wave V peak latency was seen even after the inherent delay in the ANM predictor was compensated for by delaying the predictor by 1.1 ms. It is possible that this wave V peak latency shift is due to the fact that the ANM predictor represents the signal at a stage in the auditory pathway that is closer to the wave V generator, compared to the RS predictor. This could result in a smaller delay for the wave V in the ANM TRF. However, we cannot exclude that this delay may be due to poor resolution of the TRFs after smoothing using a 2 ms Hamming window, or due to a misaligned predictor. Accounting for the inherent delay is not straightforward, as the auditory model results in varying delays based on the input stimulus properties, to better simulate realistic auditory system behavior. Our correlation method might not optimally capture this, and further work is needed to determine a more accurate method to align the ANM predictor, and to investigate the interplay between AN modelling delays and the impact of using a predictor representation that is closer to wave V generators. Furthermore, other types of auditory models could also be used to generate predictors (Dau, Püschel, & Kohlrausch, 1996b, 1996a; Osses Vecchi & Kohlrausch, 2021; Verhulst, Bharadwaj, Mehraei, Shera, & Shinn-Cunningham, 2015) and may result in improved TRFs, perhaps with clear early peaks (waves I-IV). A comparison of several auditory peripheral models in terms of simulating the auditory system is provided in (Vecchi et al., 2022). A more direct comparison of auditory model predictors for subcortical TRF estimation is provided in a recent preprint (Kulasingham et al., 2023).

### 4.4 More data required for estimating subcortical TRFs in the sound-field than with insert earphones

Incorporating the auditory nerve model reduced the amount of data needed to estimate a TRF with SNR of at least 3 dB to only 12-16 minutes. However, slightly more data was still needed to obtain clear responses (*≥* 3 dB SNR) for all participants when speech was presented in the sound-field (16 mins) compared to when speech was presented through insert earphones (12 mins). This is likely due to room acoustics affecting speech propagating through the sound-field, resulting in differences between the employed predictors and the signal reaching the ear drum, and could be alleviated by basing predictors on sound recorded close to the ear. The TRF methods used in this work closely follow the techniques provided in previous investigations into subcortical responses to speech (Bachmann et al., 2019, 2021; Maddox & Lee, 2018; Polonenko & Maddox, 2021). However, there are several possible alternatives that could be explored that may optimize the estimation of subcortical TRFs, and reduce the amount of data required for clear subcortical responses. In this work, the Cz channel referenced to the linked mastoids was used for all analysis, but other EEG channels and reference schemes have also been used for ABR ERPs (Skoe & Kraus, 2010). Indeed a multi-channel approach could allow for the incorporation of more advanced preprocessing and artifact removal methods using spatial filters such as ICA similar to what is commonly done for cortical ERPs or TRFs, albeit at the expense of increased sensors and more experimental burden. Another large difference between our method of estimating subcortical TRFs and the widely used *cortical* TRF methods is the lack of regularization. Almost all methods for estimating cortical TRFs use some form of regularization (Alickovic et al., 2019; Brodbeck et al., 2021; Crosse et al., 2021; Kulasingham & Simon, 2023), in order to reduce noise and produce interpretable TRF peaks. However, optimizing the regularization parameter could greatly increase the analysis time and the finetuning required for reasonable TRF results. We found no need to use regularization, since our goal in this work was to detect subcortical TRFs using simpler algorithms that are consistent with methods used in prior work without regularization (Maddox & Lee, 2018; Shan et al., 2023). Indeed, our work shows that it is possible to detect subcortical wave Vs using a simple artifact rejection procedure and a single-channel electrode configuration using unregularized TRFs and the ANM predictor. However, investigating more complex alternatives might provide more insights using less EEG data, albeit with larger computational costs.

### 4.5 Potential audiological applications

Clear subcortical responses to continuous speech presented in the sound-field were found for all participants on an individual level, which is crucial for potential audiological applications. Measuring subcortical neural responses to naturalistic sound-field stimuli offers the possibility of objective hearing assessment in a more life-like setting, with realistic testing conditions and using an ecologically relevant, complex stimuli, which might ultimately enable gaining a better representation of people’s hearing ability in daily life. Objective hearing testing is of special relevance for populations such as newborns or people with neurodegenerative diseases who are unable to provide reliable feedback required for behavioral hearing testing (e.g. pure-tone thresholds). Especially in these populations, eliminating the need for headphones might alleviate agitation associated with the testing situation. Measuring in the sound-field additionally offers the option to include assistive hearing devices, and investigate their effect on subcortical neural processing. For example, the effect of directionality algorithms for noise suppression in hearing aids could be evaluated at the brainstem level, similar to recent work investigating such effects at the cortical level (Alickovic et al., 2020, 2021; Andersen et al., 2021). Even though we demonstrate that subcortical responses can be obtained to speech presented in the sound-field, our work shows that such responses may have been affected by room acoustics, and points towards the possibility of improved brainstem responses when estimated using sound recorded close to the ear. Combined with the information it may reveal about hearing status, this might pave the way towards smart assistive hearing technologies in the future.

## 5 Conclusion

Our work demonstrates that brainstem responses can be detected to continuous speech presented in the sound-field, which largely corresponded to those measured when presenting a click, and when speech was delivered via insert earphones. This brings the assessment of early objective sound processing closer to real-life conditions. We show that incorporating models of nonlinear neural processing along the auditory pathway up to the brainstem improves subcortical response estimation beyond simple halfwave rectification techniques, pointing towards the importance of predictors closely matched with the neural representation of sound at the processing stage under investigation. Comparing TRFs obtained using different auditory models in predictor estimation might provide valuable experimental feedback at the brainstem level for speech processing models. Furthermore, using tailored predictors might offer the possibility to distinctly target certain processing stages along the auditory pathway. Due to effects of room acoustics, response amplitudes were reduced when sound was presented in the sound-field instead of via insert earphones, and thus slightly longer EEG recording time was needed in the sound-field. Even so, our approach measuring subcortical neural responses to continuous speech presented in the soundfield and analyzed incorporating an auditory nerve model yields clear neural responses when computed on a 16-minute portion of the data. Crucially, clear wave V peaks were obtained for each participant, which is an essential prerequisite for potential clinical applications. These insights could pave the way towards objective hearing evaluation using ecologically relevant stimuli in a more realistic setting, and future smart hearing solutions.

## Acknowledgments

This work was supported by the William Demant Foundation. The authors are also grateful to all participants for their participation in this study.

