## Supplementary Figures for "Extending Subcortical EEG Responses to Continuous Speech to the Sound-Field"

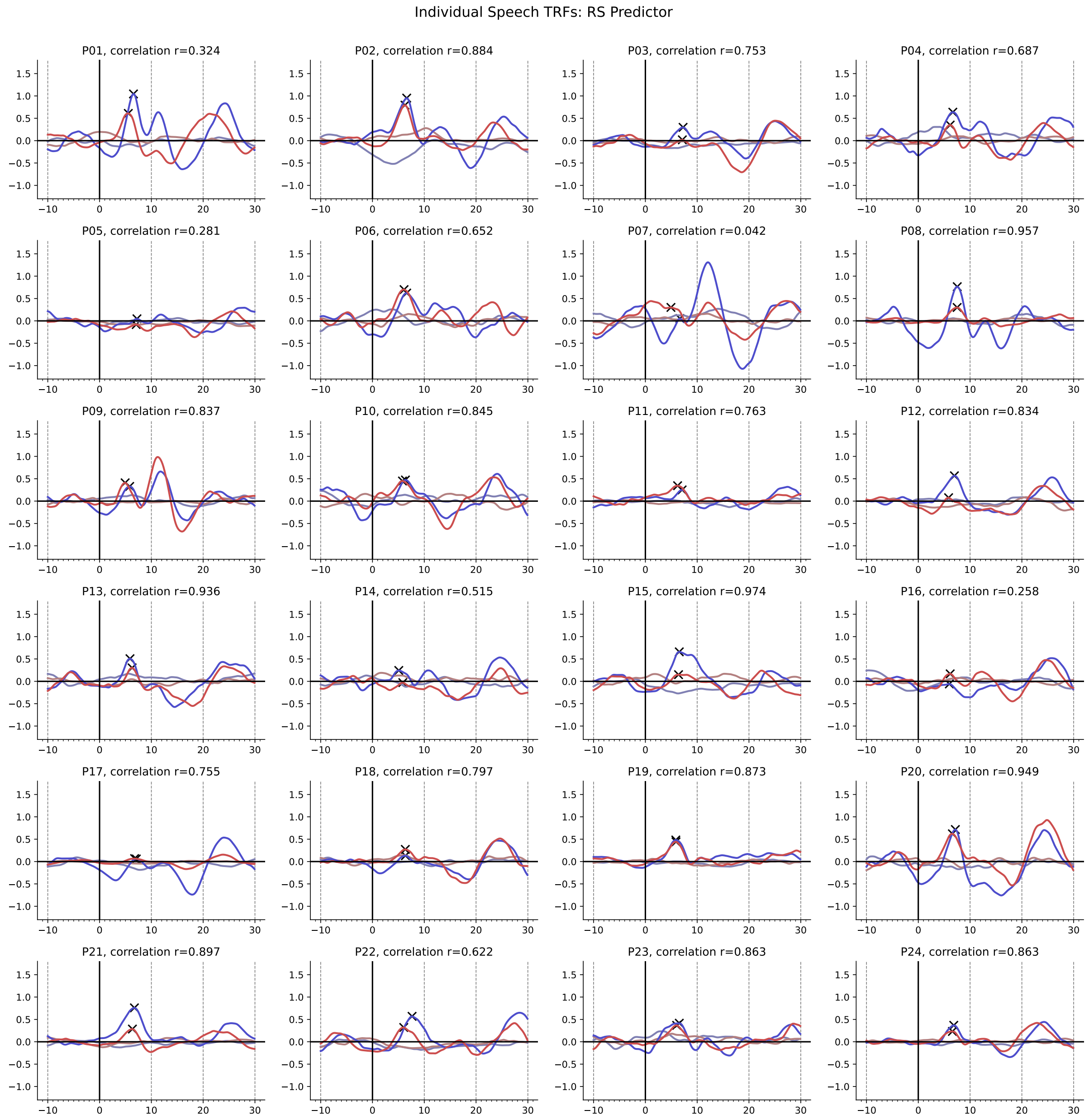


Figure S1: **Individual speech TRFs for the RS predictor.** Individual participant TRFs are shown for both insert earphone (blue) and sound-field (red) conditions, along with corresponding null models (lighter colors). In some participants, clear wave V peaks could not be determined, and the TRFs for the two conditions show substantial differences (Pearson correlation between the TRF waveforms of both conditions are shown on top of each subplot).

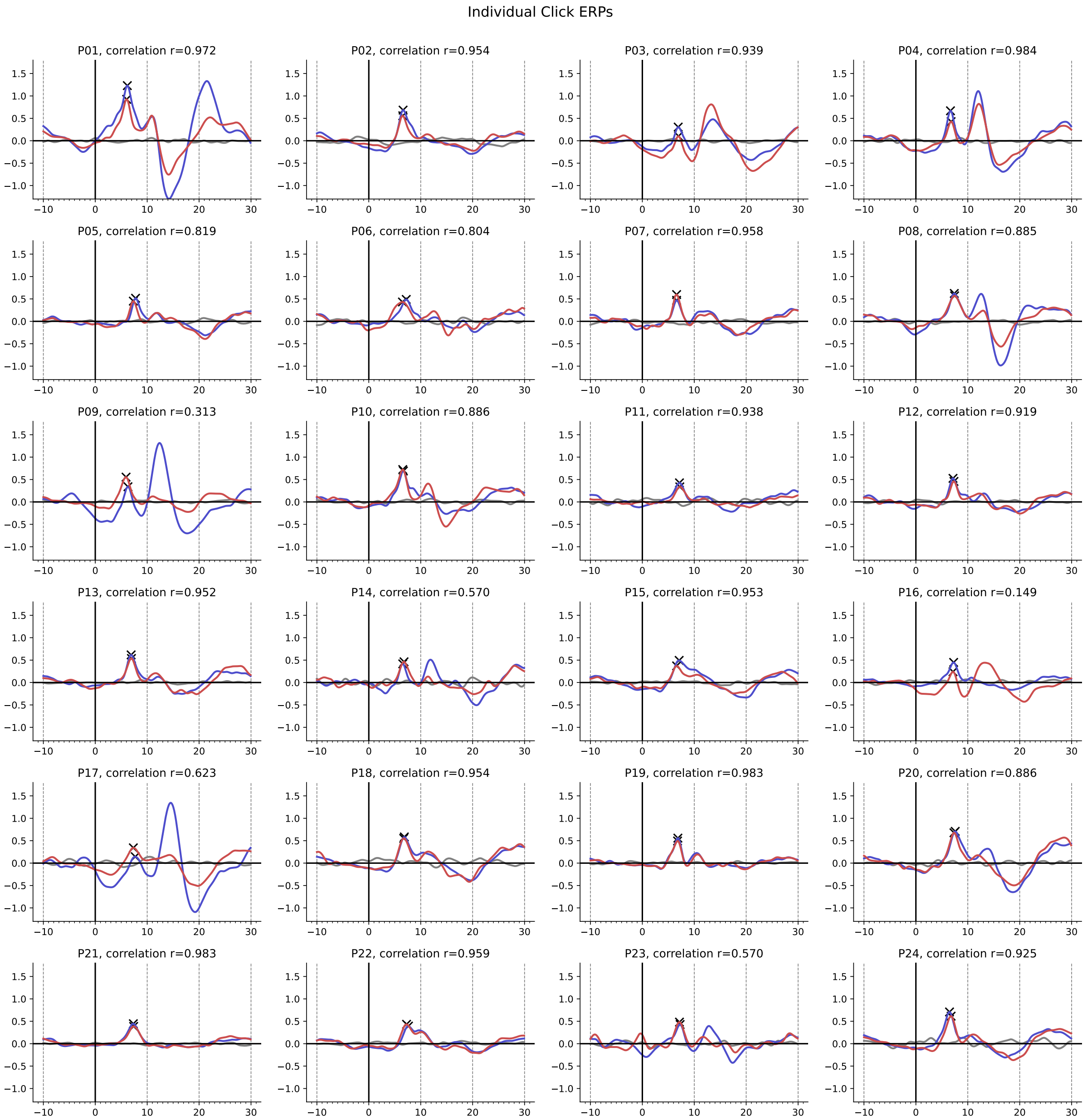


Figure S2: **Individual click ERPs.** Individual participant ERPs to clicks are shown for insert earphone (blue), sound-field (red), and control (grey) conditions. In the control condition, the eartips were not connected, and clicks were inaudible despite being presented through correctly positioned insert earphones. Clear wave V peaks are obtained in all participants and for most, the ERPs for insert and sound-field conditions are similar (Pearson correlation between the ERP waveforms of insert and sound-field conditions are shown on top of each subplot).
